# A Boolean Model of the Formation of Tumour Associated Macrophages in an *in-vitro* Model of Chronic Lymphocytic Leukaemia

**DOI:** 10.1101/2020.10.14.337642

**Authors:** Malvina Marku, Flavien Raynal, Nina Verstraete, Marcin Domagala, Miguel Madrid-Mencía, Mary Poupot, Jean-Jacques Fournié, Loïc Ysebaert, Vera Pancaldi

## Abstract

The tumour microenvironment is the collection of cells in and surrounding cancer cells in a tumour including a variety of immune cells, especially neutrophils and monocyte-derived macrophages. In a tumour setting, macrophages encompass a spectrum between a tumour-suppressive (M1) or tumour-promoting (M2) state. The biology of macrophages found in tumours (Tumour Associated Macrophages) remains unclear, but understanding their impact on tumour progression is highly important. In this paper, we perform a comprehensive analysis of a macrophage polarization network, following two lines of enquiry: *(i)* we reconstruct the macrophage polarization network based on literature, extending it to include important stimuli in a tumour setting, and *(ii)* we build a dynamical model able to reproduce macrophage polarization in the presence of different stimuli, including the contact with cancer cells. Our simulations recapitulate the documented macrophage phenotypes and their dependencies on specific receptors and transcription factors, while also elucidating the formation of a special type of tumour associated macrophages in an *in-vitro* model of chronic lymphocytic leukaemia. This model constitutes the first step towards elucidating the cross-talk between immune and cancer cells inside tumours, with the ultimate goal of identifying new therapeutic targets that could control the formation of tumour associated macrophages in patients.

## 1. Introduction

As all living cells, macrophages perceive and respond to intra- and extracellular signals in order to maintain their functions (endocytic, phagocytic and secretory, for example) by displaying a wide spectrum of specific phenotypes (polarizations) in different inducer environments. Based on their activity and the expression of specific proteins, markers and chemokines, two major subsets of macrophages have been identified, namely classically activated macrophages (M1) exhibiting a pro-inflammatory response, and alternatively activated macrophages (M2, themselves subdivided into 4 subclasses: M2a, M2b, M2c, M2d [1–3]) exhibiting an anti-inflammatory response. Additionally, multiple studies support the idea that M1 and M2 macrophages represent, in fact, the extremes of a continuous polarization spectrum of cells deriving from the differentiation of monocytes[4]. Macrophages have a plastic gene expression profile that is determined by the type, concentration and duration of exposure to the polarization stimuli in an inflammatory environment [3,5–8].

Macrophages are also found inside tumours, as part of the tumour micro-environment (TME), a complex ecology of cells that are found surrounding cancer cells including also other immune cells such as lymphocytes and neutrophils and other normal cells. In many tumours, infiltrated macrophages display mostly an M2-like phenotype, which provides an immunosuppressive microenvironment. In cancer, these *tumour associated macrophages (TAMs)* secrete several cytokines, chemokines and proteins which promote tumour angiogenesis, growth and metastasis [9–12]. Interestingly, it has been observed that, in established tumours, signals originating from cancer cells can cause phenotypic shifts in macrophages, leading to alternative functions that do not correspond to either M1 or M2 phenotypes [13]. Several studies have demonstrated that TAMs directly suppress CD8^+^ T cell activation *in-vitro* [14–17]. Mechanisms that orchestrate this process, either directly or indirectly, remain unclear [18] and warrant further exploration due to macrophages’ important impact on tumour progression.

In any given environment, the cellular processes that determine a cell’s phenotype consist in a cascade of interactions, which can be represented as a *regulatory network*, in which nodes represent proteins, enzymes, chemokines, etc., while the connections represent the type (activation or inhibition) and direction of interactions of different types (transcriptional and post-translational activations). Network modelling has found numerous applications in studying the structure and dynamic behaviour of different biological systems in response to environmental stimuli and internal perturbations [19–22]. Several computational models of different pathways involved in the inflammatory immune response have been previously published, such as: continuous, logical and multi-scale model of T cell differentiation [23–25], logical models of macrophage differentiation in pro- and anti-inflammatory conditions [26], multi-scale models of innate immune response in tumoural conditions [27], etc. An important computational model of macrophage polarization was able to detect 4 different M2 subgroups of macrophages, as a result of various combinations of pro- and anti-inflammatory extra-cellular signals [26], using exclusively literature-based knowledge on the intra-cellular regulatory interactions and pathways involved in the polarization process. Nevertheless, many important questions remain to be explored regarding the polarization states, especially in a tumour setting. More specifically, it is important to identify the pathways involved in TAM formation and to understand to what extent the macrophage plasticity facilitates this process in a TME. On the other hand, despite the wealth of quantitative information from bulk and single-cell sequencing datasets, the inference of regulatory networks based on experimental data remains a difficult challenge, with most approaches proposing a combination of both literature- and data-driven methods [28–30].

In Chronic Lymphocytic Leukemia (CLL), a B-cell malignancy in which patients accumulate large quantities of malignant CLL cells in their lymph nodes, an interesting ecology of cancer cells and immune cells is established. CLL cells are able to educate surrounding monocytes, through direct contact and cytokine signals, turning them into TAMs, which in this disease are referred to as Nurse Like Cells (NLCs) [31]. NLCs are derived from CD14^+^ monocytes and are characterised by a distinct set of antigens (CD14lo, CD68hi, CD11b, CD163hi) [32,33]. Moreover, NLCs express stromal-derived-factor-1alpha, a chemokine which promotes chemotaxis and activates mitogen activated protein kinases, ultimately leading to more aggressive cancers and better survival of these cells *in-vitro*. Through direct contact, the NLCs are able to protect the cancer CLL cells from apoptotic signals, and stimulate environment mediated drug resistance. Interactions between NLCs and CLL cells appear to be mediated by the B cell receptor, which, when stimulated, activates production of CCL3/4, initiating the recruitment of other cells, including CD4^+^ T cells and more NLCs. Another pathway that has been associated with NLCs and TAMs more in general is that of CSF-1 (MCSF). Patients with high expression of this factor usually show faster CLL progression and this gene was implicated in the production of NLCs. Also the more M1-or M2-like profile of NLCs in specific patients correlates with active and controlled disease, respectively. Analyses of the transcriptomic profile of NLCs suggest their high similarity to the macrophage M2 profile described in solid tumours, which makes studying the formation of NLCs all the more relevant in the quest of controlling TAMs in other malignancies.

NLC formation can be studied through an *in-vitro* system in which heterologous co-cultures of healthy monocytes and patient-derived CLL cells can be established to produce NLCs in absence of any other cell type. This system is particularly suited to mathematical modelling, as experimental conditions are well controlled controlled and the cell types present are limited to monocytes/macrophages and cancer cells, without the confounding effects of immune or other healthy cells.

Boolean models are discrete dynamical models, in which each component (gene, transcription factor, chemokine, cytokine, receptor, etc.) is associated with a discrete (binary) variable, representing its concentration, activity or expression. Despite the complex processes relating the transcription of a gene into an mRNA and its subsequent translation into a protein with possibly post-translational modifications, in this paper we consider a single node for gene, mRNA and protein, such that a link between two transcription factors signifies that one of them affects transcription of the gene of the other. The future states of each component are determined by the current states of its regulators, as given by a Boolean function that represents the regulatory relationships between the components according to the logic operators AND, OR and NOT. The *state* of the system at each time point is given by a binary vector, in which each element represents the state of the corresponding component (ON/OFF) [23,34]. Starting from an initial state, as time passes the system will follow a trajectory of states reaching one of many attractors that can be a single stable state (fixed point) or a set of recurrent states (limit cycle). Attractors usually represent specific phenotypes, such as cellular differentiated states, cell cycle states, etc. Despite their coarse-grained description, Boolean models have been successfully used to capture real-world biological features like, for example, the mechanisms of cell fate decision [35], hierarchical differentiation of myeloid progenitors [36], dynamical modelling of oncogenic signalling [37], amongst many other applications [38–40]. One of their main advantages is the simplicity of performing *in-silico* experiments simulating a variety of mutant and knockout conditions, and the possibility of obtaining qualitative or semi-quantitative results without requiring experimentally-derived parameter values, as needed by differential equations. Starting from a pathway diagram describing a biological process, and adding logic rules, Boolean models allow us to model the process, uncover the main regulators, and run simulations.

Understanding the mechanisms of TAM formation is of particular interest because of their pro-tumoural activity which hampers T cell cytotoxic activity. In this study, we therefore follow two lines of enquiry: (i) we reconstruct a macrophage polarization regulatory network using literature and extend it based on transcriptomic data from an *in-vitro* model of NLC formation, (ii) we implement a Boolean model of monocyte differentiation into NLC simulating these *in-vitro* cultures.

## 2. Results

### 2.1. Reconstruction of the regulatory network leading to NLC formation

To reconstruct the gene regulatory network (GRN) governing the formation of NLCs, we started from a previous macrophage polarization GRN [26] and extended it in order to include specific extracellular signals found in the Chronic Lymphocytic Leukaemia (CLL) context and other intra-cellular components involved in NLC formation. The network extension was based on extensive literature review and transcription factor (TF) activities estimation for each phenotype. Briefly, we used transcriptomics data for monocytes, M1, M2 and NLCs to calculate the TF activities in each condition, and chose the TF with the highest activities in each phenotype (see Methods, Section 5.3). For NLCs, we identified specific TFs using a set of 17 microarray expression profiles [33], which interestingly have higher activities than in M1 and M2. Particularly, HMGB1 and HIF1 are linked to the pro-tumoural activity of NLC (Appendix), and were considered as key regulators that determine the distinct phenotypes between M2 and NLC.

The main characteristics of these 3 types of macrophages are given in Table 1. A short description of the profiles for the main macrophage phenotypes is given in Appendix, however, a detailed explanation of the mechanisms, pathways and components involved in the polarization process can be found in the cited papers and the references therein.

**Table 1.** The main charecteristics of M1, M2 and NLC phenotypes according to (i) activators, (ii) secreted cytokines or expressed genes, and (iii) functions in tumoural environments (see Appendix).

|  | **M1** | **M2** | **NLC** |
| --- | --- | --- | --- |
| **Activated by** | IFNγ<br>LPS | Immune complexes<br>IL-4<br>IL-13<br>IL-10 | IL-10<br>TGF-β<br>CSF-1 |
| **Secrete** | Th1 inducing cytokines<br>TNFα<br>IL-12<br>IL-18<br>IFN$\alpha/\beta$<br>IL-1β<br>IL-6 | IL-10<br>TGF-β<br>VEGF<br>EGF | IL-10<br>TGF-β<br>IL-1Ra<br>HIF1/2<br>PD-L1; B7-H4;<br>TNFSF13/B;<br>VEGF<br>chemokines: CXCL12, CXCL13 |
| **Function** | **Anti-tumoural activity:**<br>releasing nitric oxide (NO);<br>presenting tumour antigens to CD4+ Th1 cells;<br>driving the activity of cytotoxic CD8+ T cells at the tumour site | **Anti-inflammatory processes:**<br>Th2 responses (M2a);<br>Downregulation of immune response (M2b);<br>matrix deposition and tissue remodelling (M2c) | **Promote tumour growth:**<br>secretion of soluble immunosuppressive agents;<br>expression of contact-dependent immunosuppressive receptors (PD-L1, B7-H4) leading to enhancing CD8+ T cell infiltration<br>high levels of HIF1 and HIF2 which leads to expression of genes associated with pro-tumoural activity |
| **References** | [1,41–43] | [1–3,43] | [12,13,17,42,44] |

The inferred regulatory network of macrophage polarization is given in Figure 1. It contains 10 extracellular signals, 30 intra-cellular components, most of them being TFs and interleukins, and 3 outputs, which are used as readouts, namely M1 polarization, M2 polarization and NLC. Pathway enrichment analysis [45] showed that most of the components are involved in the JAK-STAT signalling pathway, pathways related to cancer, Th17 cell differentiation, cytokine receptor interaction and other inflammatory conditions.

**Figure 1.**
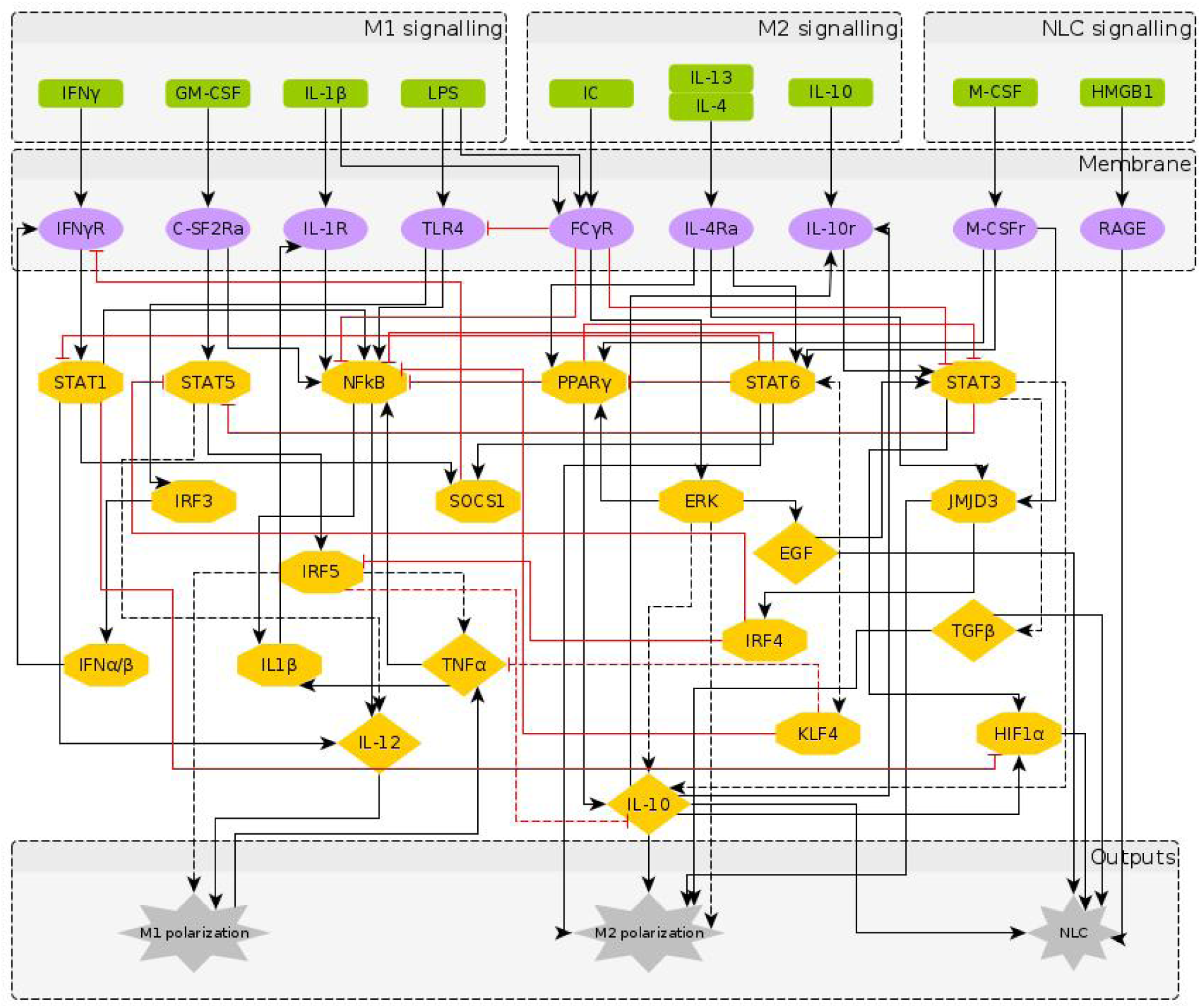
The regulatory network of macrophage polarization: Nodes in green represent the extra-cellular signals, classified as M1, M2 and NLC inducers; nodes in purple represent receptors in the macrophage membrane, usually activated upon contact with cells in the outer environment; nodes in yellow represent the transcription factors and chemokines involved in the polarization process, as an intermediate step or as an output. The interactions between components can be either activation (black) or inhibition (red). The dashed arrows indicate indirect effects, in which the targets are the end-products, i.e. intermediate interactions are involved but not represented in the network.

### 2.2. A Boolean model of macrophage polarization

Starting from the regulatory network in Figure 1, the Boolean functions for each component are given in Table 2. Here, the Boolean functions were based on published experimental evidence from the literature. The numerical simulations were performed considering all the possible initial intracellular conditions and combinations of stimuli, while applying the synchronous updating method to calculate the system’s attractors (section 5.1). The simulation results show that the system reaches 1384 fixed point attractors, while other cyclic attractors of length 2 and 3 were also present. For our scope, in the following paragraphs we focus only on the fixed point attractors. It is important to note that fixed point attractors are time invariant, i.e. the number of fixed points is not affected from the updating method chosen, while the number of cyclic attractors and their characterisitcs (period, basin of attraction) depend on the updating method (section 5.1). Here and throughout the paper, we will refer to an attractor as the binarized expression profile which we assign to a polarization state (or a phenotype). To attribute the attractors to certain phenotype categories, we removed all the input nodes (extracellular signals) from the attractors, thus reducing the attractors’ space to 214 fixed points.

**Table 2.**
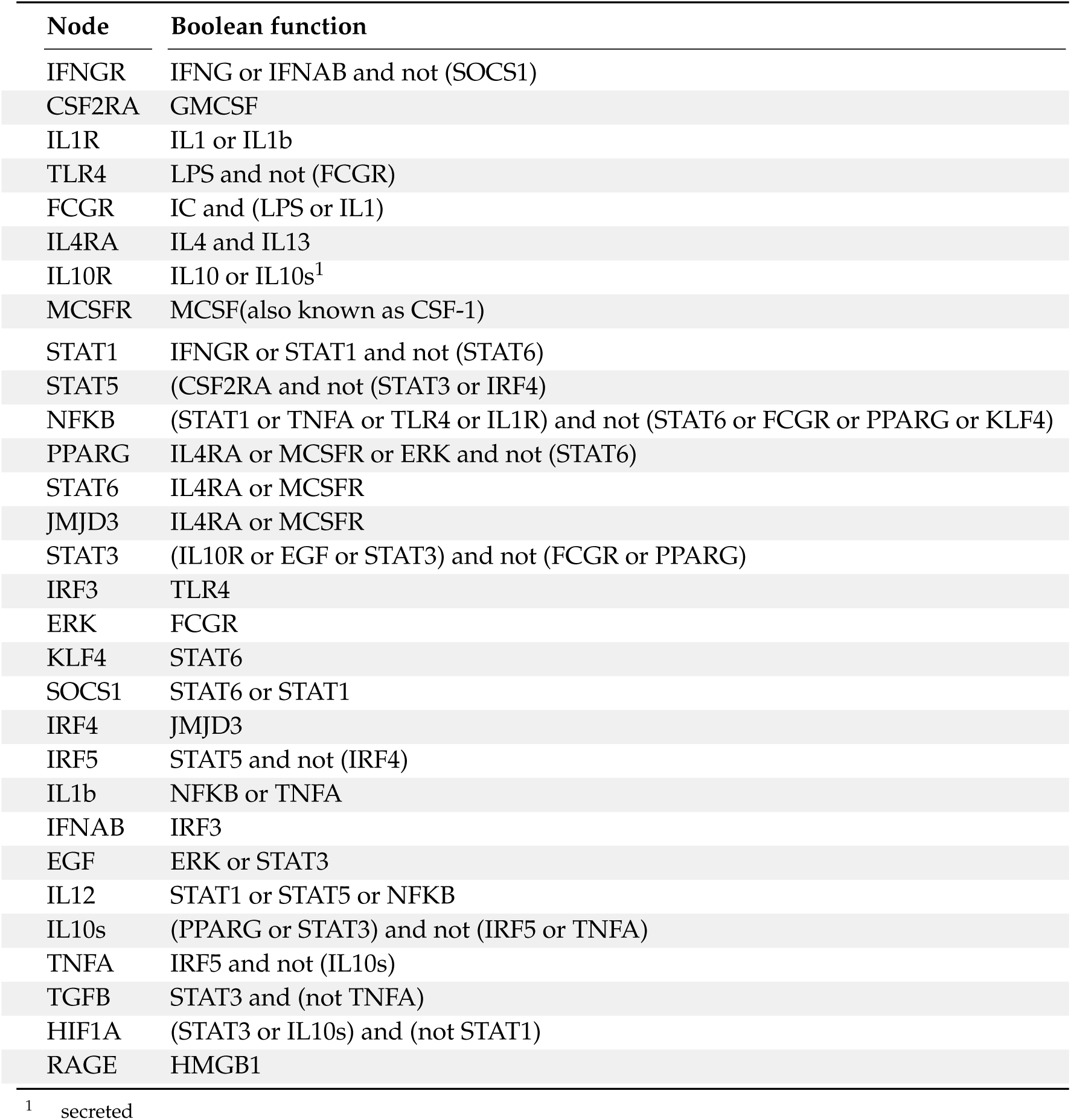
Boolean rules of the 30 intra-cellular nodes of the macrophage polarization network

| Node | Boolean function |
| --- | --- |
| IFNGR | IFNG or IFNAB and not (SOCS1) |
| CSF2RA | GMCSF |
| IL1R | IL1 or IL1b |
| TLR4 | LPS and not (FCGR) |
| FCGR | IC and (LPS or IL1) |
| IL4RA | IL4 and IL13 |
| IL10R | IL10 or IL10s<sup>1</sup> |
| MCSFR | MCSF(also known as CSF-1) |

<sup>1</sup> secreted

| Node | Boolean function |
| --- | --- |
| STAT1 | IFNGR or STAT1 and not (STAT6) |
| STAT5 | (CSF2RA and not (STAT3 or IRF4) |
| NFKB | (STAT1 or TNFA or TLR4 or IL1R) and not (STAT6 or FCGR or PPARG or KLF4) |
| PPARG | IL4RA or MCSFR or ERK and not (STAT6) |
| STAT6 | IL4RA or MCSFR |
| JMJD3 | IL4RA or MCSFR |
| STAT3 | (IL10R or EGF or STAT3) and not (FCGR or PPARG) |
| IRF3 | TLR4 |
| ERK | FCGR |
| KLF4 | STAT6 |
| SOCS1 | STAT6 or STAT1 |
| IRF4 | JMJD3 |
| IRF5 | STAT5 and not (IRF4) |
| IL1b | NFKB or TNFA |
| IFNAB | IRF3 |
| EGF | ERK or STAT3 |
| IL12 | STAT1 or STAT5 or NFKB |
| IL10s | (PPARG or STAT3) and not (IRF5 or TNFA) |
| TNFA | IRF5 and not (IL10s) |
| TGFB | STAT3 and (not TNFA) |
| HIF1A | (STAT3 or IL10s) and (not STAT1) |
| RAGE | HMGB1 |

### 2.3. Phenotype identification through interpretation of the attractors

The large attractors’ space raises the challenge of interpreting its biological meaning. To categorize the attractors in specific polarization states, two different methods were used: 1) a supervised literature-based method using the expression profiles of the macrophage phenotypes taken from the literature, and 2) an unsupervised method grouping attractors based on their similarity and then applying clustering algorithms to assign them to specific phenotypes.

#### 2.3.1. Intepreting attractors based on a supervised method

To identify the main phenotypes detected by the model, we categorized all the attractors according to the expression profiles of M1, M2 and NLC known from the literature, as described in Table 1 and Appendix:

- M1: IL-12, NF-*κ*B, TNF*α* and STAT1 or STAT5 active;
- M2: IL-10, STAT3 or STAT6, PPAR*γ* active;
- NLC: TGF*β*, HIF1*α*, EGF, RAGE active;
- M0: M0 attractors + Attractors not falling in any of the above categories.

It is important to note that the M1, M2 and NLC categories were considered as mutually exclusive; therefore the rest of the attractors were categorized together with M0, in a cateogry apart that includes all the attractors exhibiting characteristics of both M1 and M2 phenotypes, or corresponding to states without biological significance. Interestingly, we found that most of the attractors fall into the M2 (*≈*67.3%) category, followed by the M1 (*≈*4.7%) category and NLC (*≈*2%) subset (Figure 2), indicating the high likelihood for the system to reach one of the anti-inflammatory polarization states. The similarities between attractors falling in each category were estimated by calculating the Jaccard-Needham distances (dist_values *∈* [0, 0.5]). Considering the low values of binary distances between attractors in each category, we then calculated the average attractor states (Figure 2 (b)-(e)). Importantly, we observe that these averaged attractors largely correspond to the expected expression profiles for M1, M2 and NLC defined above. A principal component analysis shows the main identified clusters of attractors corresponding to each phenotype (Figure 3). From the plot, we can easily observe that NLC attractors are not well separated from M0, which can be explained considering that a large number of attractors in our M0 category have profiles intermediate between M1 and M2 and NLCs are also thought to have an intermediate profile.

**Figure 2.**
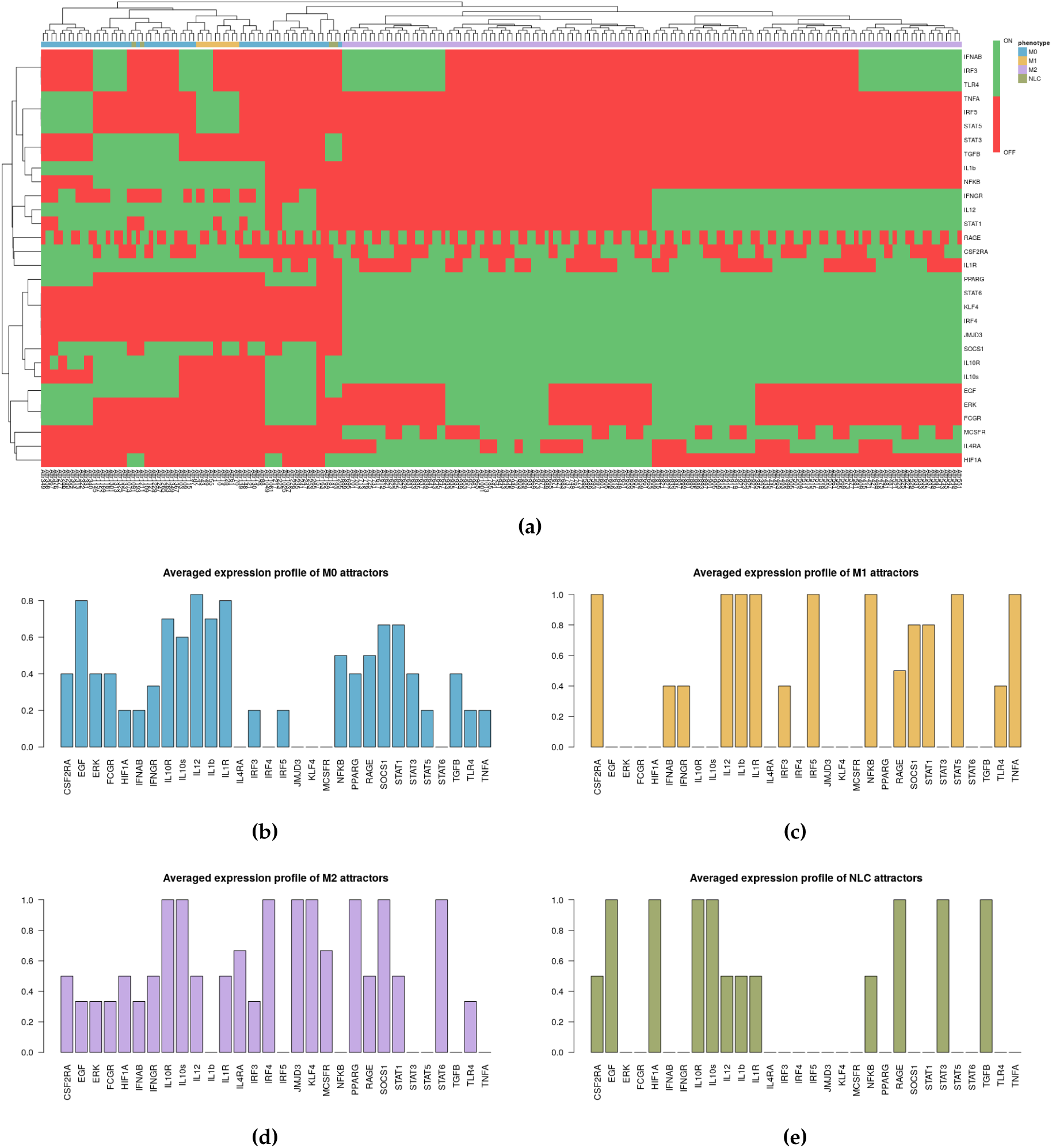
(a) Heatmap of 214 attractors. (b)-(e) Averaged attractors for each category: M0, M1, M2 and NLC.

**Figure 3.**
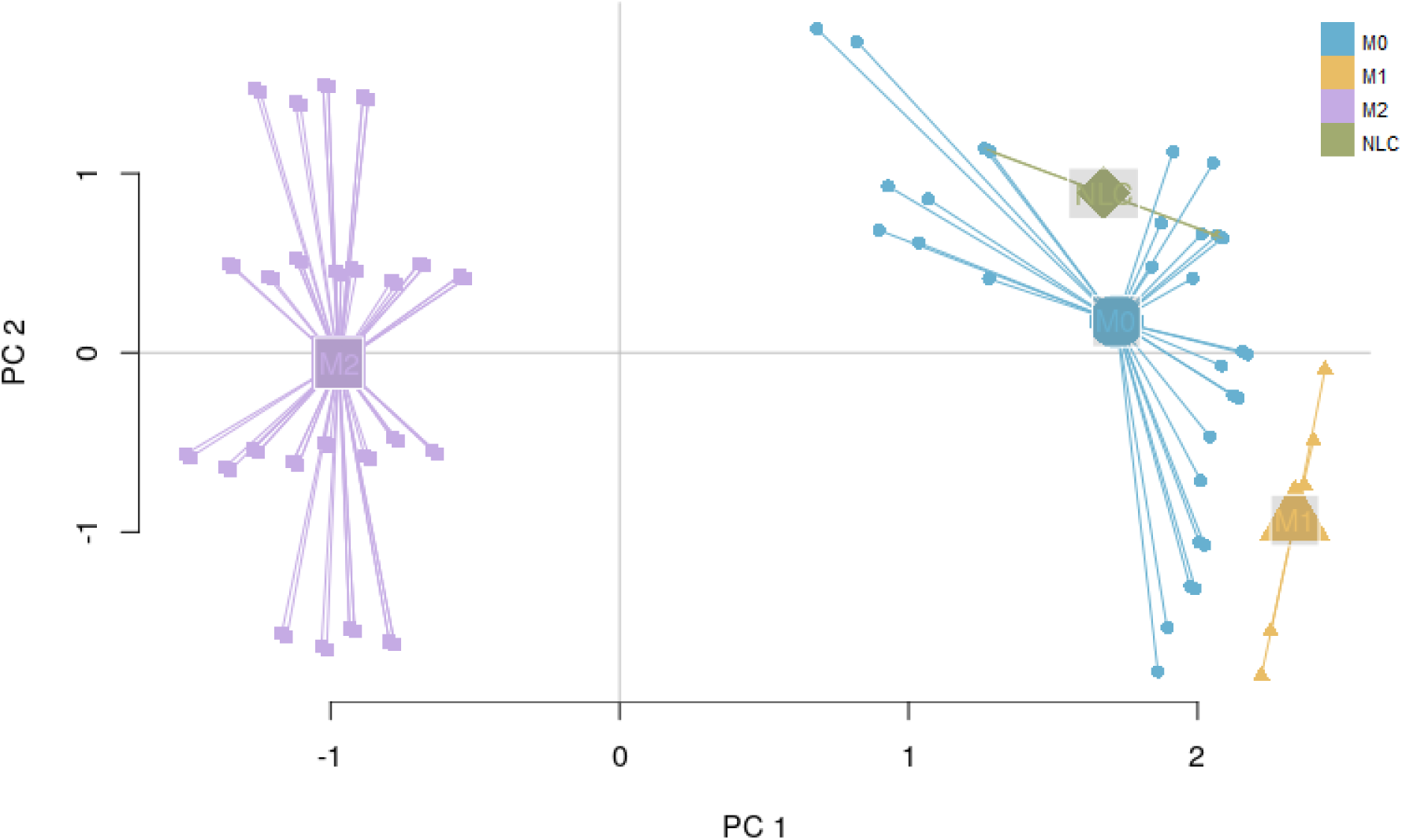
PCA of 214 attractors: M1 and M2 attractors are observed in distinct clusters, while NLC attractors appear in between the two extremes of the polarization spectrum.

#### 2.3.2. Interpreting attractors based on an unsupervised method

Alongside with the supervised method, we also performed unsupervised clustering on the attractor space, in order to investigate whether the main phenotypes we expect in this system can be recovered in an unbiased way just exploring the structure of the attractors’ space. We hypothesise that the attractors corresponding to the same phenotype category will be characterized by a small binary distance and consequently will fall into the same cluster. To this end, we first estimated the similarity among the attractors by calculating the Jaccard-Needham distance [46]. We then applied hierarchical density based clustering on the Jaccard-Needham distances (Figure 4) to identify the main attractor clusters. As can be seen from the heatmap, 5 main clusters are detected: one of them (Cluster 4) corresponds to the zero-attractors (attractor 1: all the components in OFF state, attractor 2: all the components in OFF state, except from *expr*_*RAGE*_ = 1) and it was not considered for further analysis. A closer look at the averages of the attractors falling in each cluster highlights the detected expression profiles (Figure 4 (b)-(e)). Based on the averaged expression profiles of attractors in each cluster, we observe a clear representation of M1, M2 and NLC phenotypes, respectively Cluster 5 *→* M1: IL-12, IL-1R, NF-*κ*B, STAT1, TNF*α* highly expressed, Cluster 2 *→* M2: IL-10, IL-10R, JMJD3, KLF4, IRF4, PPAR*γ* and STAT6 highly expressed, and Cluster 3 *→* NLC: EGF, HIF1*α*, RAGE, TGF*β* and IL-10 highly expressed. Considering the high expression of both M1, M2 and NLC components, we attribute Cluster 1 to M0.

**Figure 4.**
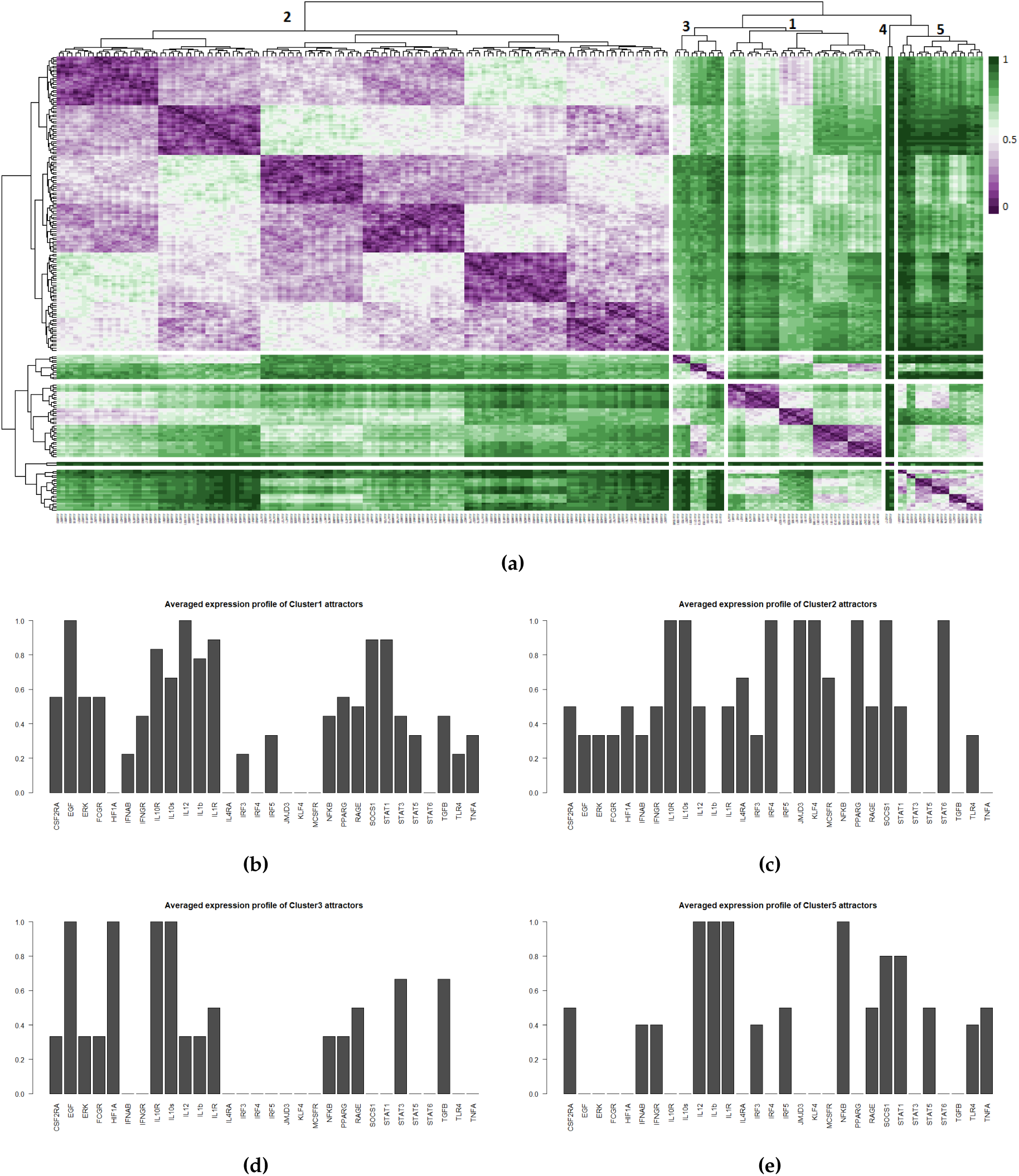
(a) Heatmap of Jaccard-Needham distances of 214 attractors: 5 main clusters can be observed. Cluster 4 contains the attractor with all nodes in the OFF state and was not considered for further analysis. (b)-(e) Averaged attractors for each cluster in (a).

#### 2.3.3. Robustness of attractor interpretation independent of annotation method

While choosing between supervised and unsupervised methods one must consider some advantages and disadvantages. Supervised approaches can ensure a specific match between the observed attractors and prior biological knowledge of each phenotype, which can be an issue when the attractors can correspond to uncharacterised biological states and can be limited to the use of existing knowledge. On the other hand, unsupervised methods offer the simplicity of detecting the different state categories in a more unbiased way and possibly to identify unknown intermediate phenotypes in the polarization spectrum of the macrophages.

For a more quantitative comparison between the supervised and the unsupervised methods, we calculated the Pearson correlation coefficient between the averaged expression profiles obtained from each phenotype and each cluster (Figure 5). Our results show the accuracy of the unsupervised method in capturing the M1 (*corr*_*coe f f* = 0.92), M2 (*corr*_*coe f f* = 1) and NLC (*corr*_*coe f f* = 0.91) phenotypes, while the M0 category matches best with Cluster 1 with *corr*_*coe f f* = 0.97, not corresponding to any phenotype.

**Figure 5.**
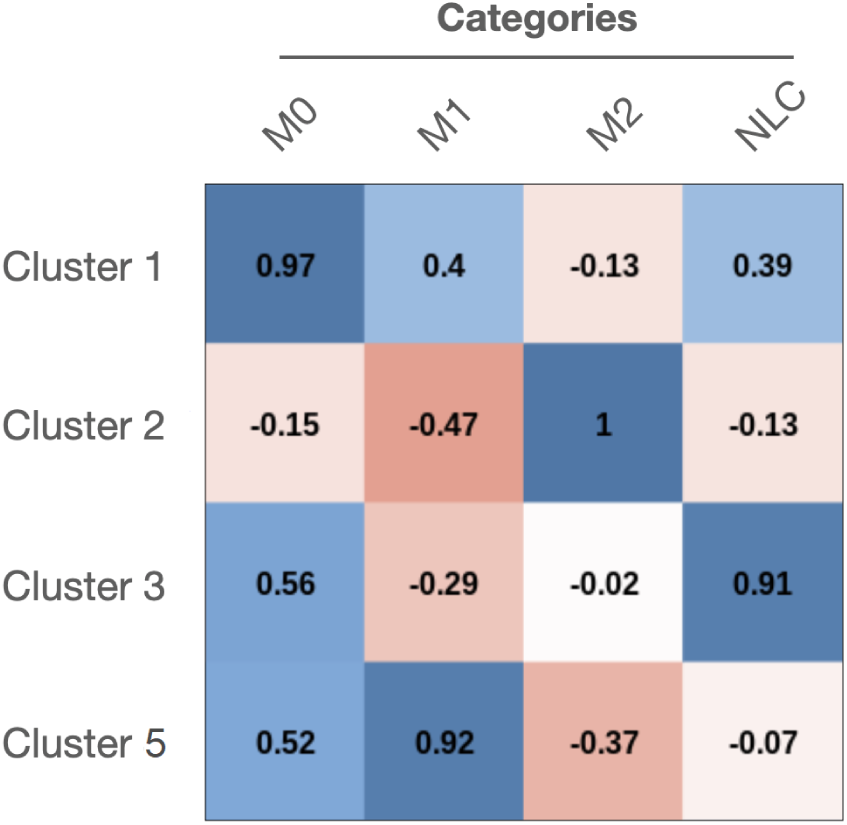
Matrix of Pearson correlation between M0, M1, M2 and NLC categories and the 4 biologically relevant clusters.

## 3. Model validation through *in-silico* perturbations

To validate the model, we performed several simulations mimicking specific environmental conditions consisting of M1, M2 or NLC signals only. Previous wet-lab experiments have shown that in co-cultures of monocytes and CLL cells, the CLL signal will elicit the differentiation of monocytes into NLCs. We studied the attractor space in the presence of only CLL signals (M-CSF and HMGB1) while considering all the possible combinations of intra-cellular signals. We then hypothesised that the presence of only a specific phenotype signal inducer (M1, M2 or NLC) would shift the macrophages polarization towards the corresponding phenotype and performed different simulations setting the signals favouring a certain phenotype to the ON state. Indeed, our simulations showed that the presence of specific signals (grouped as M1, M2 and NLC signals) would activate certain pathways that subsequently lead to the corresponding polarization state. Table 3 recapitulates the simulations performed by selecting only specific stimuli, the observed attractors’ categories, the expression profiles of each polarization state and the network representation of active/inactive nodes/edges under these conditions. Interestingly, we observed that while the presence of M1 and M2 signals leads to the activation of their corresponding phenotypes, NLC signals activate both M2 and NLC polarization states, which reinforces the shared pro-tumoural activity of both phenotypes in the TME.

**Table 3.**
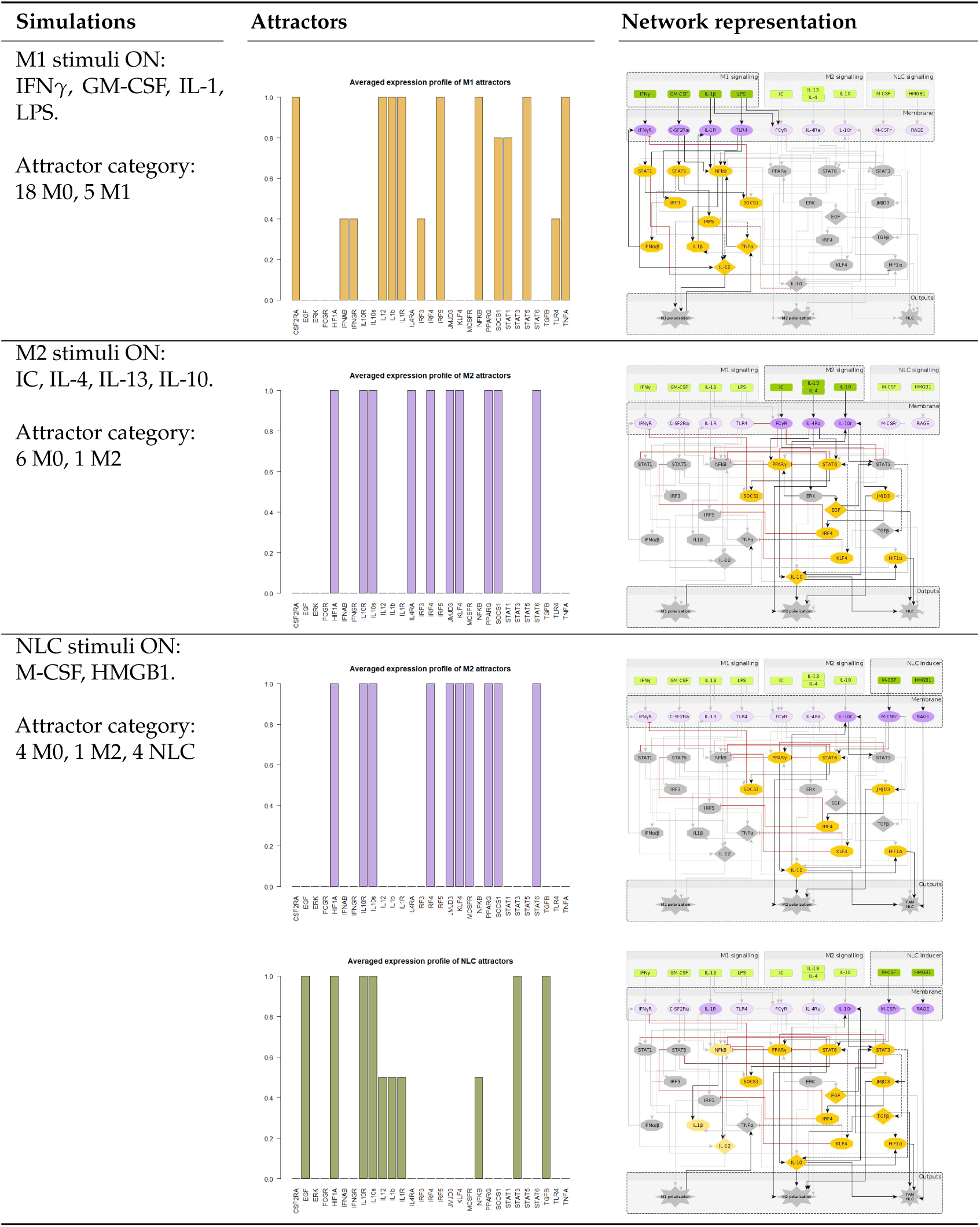
Simulations of environmental signals consisting of M1-, M2- and NLC-inducing signals only.

Additionally, several experimental studies on the effects of mutants and knock-outs on macrophage polarization states have been previously published [47–50]. Here, we performed simulations of knock-outs, as summarized in Table 4. Analysing the attractors’ space, we observed a complete loss of M2 phenotype in STAT6^*–*/*–*^, IRF4-JMJD3 axis KO and a significant decrease of M2 attractors in PPAR*γ*^*–*/*–*^ and IL-4R*α*^*–*/*–*^, a complete loss of M1 phenotype in IRF5^*–*/*–*^ and STAT5^*–*/*–*^, and a significant decrease in M1 attractors in STAT1^*–*/*–*^. Additionally, we observed a complete loss of NLC phenotype in STAT3^*–*/*–*^ and EGF^*–*/*–*^. These results show that our model recapitulates the experimental observations in mutant conditions, as well as polarization outputs in the presence of different extra-cellular signals.

**Table 4.** *In-silico* experiments with knock-outs [50].

| Knock-out | Expected effect on polarization | Model results |
| --- | --- | --- |
| STAT6 | Complete knock-out<br>Loss of M2 [51] | Complete loss of M2<br>M1 and NLC attractors not affected |
| PPARγ | Conditional knock-out<br>Loss of M2 [49] | Decrease number of M2 attractors<br>M1 and NLC attractors not affected |
| IL-4Rα | Conditional knock-out<br>Loss of M2 [52] | Decrease number of M2 attractors<br>M1 and NLC attractors not affected |
| IRF5 | Complete knock-out<br>Loss of M1 [53] | Complete loss of M1<br>M2 and NLC attractors not affected |
| STAT5 | Complete knock-out<br>Loss of M1 [54] | Complete loss of M1<br>M2 and NLC attractors not affected |
| IRF4 - JMJD3 axis | Complete knock-out<br>Loss of M2 [55] | Complete loss of M2<br>M1 and NLC attractors not affected |
| **Other simulations** |  |  |
| STAT3 | Complete loss of NLC<br>Increase of M1 attractors<br>M2 attractors not affected |  |
| EGF | Complete loss of NLC<br>M1 and M2 attractors not affected |  |
| STAT1 | Significant loss of M1<br>M2 and NLC attractors not affected |  |

## 4. Discussion

The results reviewed in the previous sections highlight the various ways in which network-based dynamic models can be used to recapitulate the known characteristics of biological systems, as well as to predict new behaviours in specific conditions. Particularly, despite their limitations to a qualitative description, Boolean models yield a comprehensive picture of a system’s dynamics, including all the attractors of the system and the effects of mutants. Here, our main focus lies in identifying the mechanisms that trigger the formation of NLCs in Chronic Lymphocytic Leukaemia, a macrophage polarization state distinct from the ones that can be obtained with monocyte *in-vitro* differentiation. Despite a large body of work on macrophage polarization, the phenotypic profile and formation of tumour associated macrophages have not been fully elucidated yet, due to the difficulty of isolating these cells from tumours. For this reason, we extend a previously published Boolean model of macrophage polarization [26], by including specific nodes (genes, transcription factors and receptors) that characterise the NLC profile. We then apply Boolean rules to the regulatory network to study the system’s asymptotic behaviour, when starting from all the possible initial conditions. The main macrophage polarization states (phenotypes) were matched to the attractors first by applying constraints on the value of specific network components (literature-based constraints) and subsequently using unsupervised clustering of the attractors according to their (binary) similarities. Importantly, the model results show that the attractor categories obtained by both supervised and unsupervised methods, qualitatively match the M1, M2 and NLC profiles, while highlighting specific characteristics of NLCs that distinguish them from M2 macrophages. In addition, the unsupervised method, although less accurate than the supervised approach in characterizing the phenotypes, was shown to correctly separate the phenotypic profiles in the absence of any constraint or previous knowledge. Clustering of attractors with more powerful techniques [56,57]) would make the unsupervised method suitable especially in Boolean modelling of large networks for which prior biological knowledge is not available.

It is important to note that both the network extension and the Boolean functions were based on extensive literature review, which raises the difficulty of literature-based network inference methods for large regulatory networks. A more data-driven approach to network inference will be considered for future work [58,59].

The ultimate test of the model presented would be to compare our *in-silico* signatures for the different attractors with experimental data measuring the state of each of our model components, possibly through transcriptomic or proteomic characterization of each cell type. However, the multiple levels at which the state of a component can be experimentally determined (gene expression, protein level, protein activation state) reduce our expectations for finding a clear match. Even for the well-characterised biological processes of macrophage polarization, all experimentally derived readouts of the different phenotypes come from the detection of proteins on cell membranes, leaving gaps in our understanding and justifying the need for data-driven approaches.

Taken together, our model can describe macrophage polarization in different environments and mutant conditions. The inflammatory and cancer environments are characterized by a complex combination of stimuli, which drive the polarization process of monocytes towards specific macrophage phenotypes. In our network, we include the most significant pro- and anti-inflammatory signals, as well as important cytokines that are involved in NLC polarization, like CSF-1 (M-CSF in our model) and HMGB1. Despite the specific characteristics of the tumour micro-environments in solid cancers compared to the *in-vitro* model considered here, we believe that common polarization pathways are also involved in the formation of tumour associated macrophages (TAMs) in solid tumours, which have so far been modeled with a stronger emphasis on the inter-cellular aspects than on the molecular details [60–62]. Further work will be needed to establish whether our model can be useful more generally in different cellular environments. Overall, we hope that our model will encourage new empirical investigations on the complex nature of cell-cell interactions in the TME and the role of TAMs in cancer prognosis and treatment.

## 5. Methods

### 5.1. Boolean model Implementation

In the Boolean model each component (gene/mRNA, protein, chemokine) is associated with a discrete (binary) variable, representing its concentration, activity or expression. Time is considered to be implicit and the future states of each component were determined by the current states of their regulators, given by a Boolean function of *m*_*i*_ = 1, 2, …, *N* regulators of component *X*_*i*_. Each Boolean function represents the regulatory relationships between the components and is expressed via Boolean operators AND, OR and NOT. The state of the system at each time point is given by a binary vector, whose *i*th element represents the state of the component *X*_*i*_ [23,34]. The set of all possible states and their transitions can be represented by a state transition graph, in which the nodes are the system’s states (represented as binary vectors) and the directed edges are the transitions between them. The exponential function between the number of components and the state space size makes the graphical representation possible for only small networks. In Boolean models time is discrete and implicit: starting from an initial state, the system will follow a trajectory of states and, because of the finite state space, it reaches an attractor (stable states or limit cycles). To evaluate the state of each node at each timestep, two main updating methods have been proposed [21,63]:

- *synchronous updating method*: at each time step, all the nodes are updated simultaneously, assuming that all the interactions in the system require the same time to occur. Importantly, the state space is characterized by non-overlapping basins of attractions.
- *asynchronous updating method*: at each time step, the updated nodes are chosen randomly (General Asynchronous, Random Asynchronous) or according to their *characteristic updating time*, while the system’s state will be characterized by overlapping basins of attractions.

It is important to note that fixed point attractors are time invariant, i.e. do not depend on the updating method. Our network is composed of *N* = 40 components and has 2^40^ possible states. Choosing the synchronous update method we obtain all transitions between them and consider the final attractors. The model was implemented using the BoolNet [48] R package [64].

### 5.2. Calculating the attractor similarity matrix

Given Ω a space of binary *N*-dimensional vectors *Z* defined as

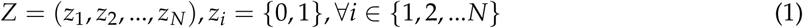

we define 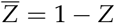 to be the complement of the binary vector *Z*. For each set of binary vectors *Z*_1_, *Z*_2_ *∈* Ω let *S*_*ij*_ be the number of occurrences of matches, with *i ∈ Z*_1_ and *j ∈ Z*_2_ being in the corresponding positions. In this way *S*_11_(*Z*_1_, *Z*_2_) = *Z*_1_ *· Z*_2_ and 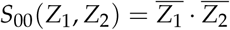. Based on *S*_*ij*_, different measures exist, to calculate the similarity/dissimilarity between two binary vectors [46]. For our purpose, we calculated the *Jaccard-Needham* measures, defined as follows:

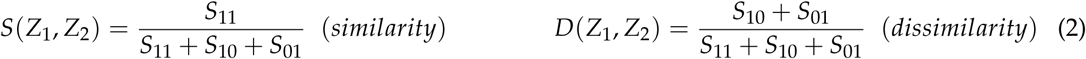

### 5.3. Calculating the transcription factor activities

Microarray data used in this publication were downloaded from the NCBI repository Gene Expression Omnibus (GEO) database. M1 and M2 Macrophages microarray data accession number is GSE5099. Our previously published NLC microarray dataset can be found under accession number GSE87813 and was processed as described in [33]. Raw microarray datasets were then normalized using the RMA (Robust Multi-arrays Average) normalization method and batch corrected. Transcription factors activities were estimated using the Dorothea R package. Dorothea is a TF-regulon interaction database giving each interaction a confidence level. Here, levels of confidence of interactions from A to E were taken into account. The VIPER algorithm was used to estimate TF activities based on Dorothea interactions and our expression data [65,66].

## Author Contributions

Conceptualization, M.M. and V.P.; methodology, M.M. and V.P.; software, M.M. and M.M.-M.; resources, M.M., N.V., M.D., M.P., J.-J.F., L.Y. and V.P.; validation, M.M. and F.R.; data curation, F.R.; original draft preparation, M.M., V.P. and N.V.; review and editing, M.D., M.P., J.-J.F. and L.Y.

## Acknowledgments

The authors thank Alexis Coullomb, Ting Xie, Julien Pernet, Maria Fernanda Senosain Ortega and Julie Bordenave for advising and critical reading of the manuscript.

## Appendix A Appendix A.1 M1 pathway

The M1-like pro-inflammatory polarization state is applied to pro-inflammatory macrophages and can be obtained upon stimulation of those cells with IFN*γ* or LPS which cause release of Th1-inducing cytokines including tumour necrosis factor *α* (TNF*α*), IL-12, IL-6, IL-1*β*, IL-18 and IFN*α*/*β* [1,41,42]. The M1 macrophages metabolism rely on oxidative glycolysis [67] and intrinsically their polarization is linked with activation of STAT1, IRF5 and NF-*κ*B [68]. M1-like macrophages are linked in fighting bacterial infections and intracellular pathogens. Additionally they show potent anti-tumoural activity which manifests mainly through: *(i)* release of large amount of nitric oxide (NO), which in turn is able to kill the cancer cells as a result of DNA damage, disruption of mitochondrial activity and limitation of iron availability, and *(ii)* presentation of tumour antigens to CD4^+^ Th1 cells and driving the activity of cytotoxic CD8^+^ T cells at the tumour site [43].

## Appendix A.2 M2 pathway

M2-like macrophages include a wide variety of phenotypes involved in resolving of the inflammation. The M2 activation can be induced by stimulation with IL-4, IL-13, immune complexes and IL-10. The anti-inflammatory and regenerative activity of M2 macrophages come from abundant release of IL-10, TGF-*β*, VEGF and EGF [1,43]. M2 macrophages depend strongly on oxidative phosphorylation [67] and the main TFs driving their polarization-state are: STAT6, PPAR*γ*/*δ*, IRF4, JMJD3 [68]. Depending on the anti-inflammatory processes M2-like macrophages are involved in, they manifest diverse phenotypes including: M2a - Th2 responses and killing and encapsulation of parasites, M2b – immunoregulation, M2c – matrix deposition and tissure remodeling [1–3].

Tumor-associated macrophages belong to the group of cells that arise upon the contact with cancer cells and tumor microenvironment (TME). They can show characteristics of both M1 and M2 state, nevertheless upon prolonged presence in the TME the M2 characteristic becomes prevalent. TAMs influence the properties and dynamics of TME, although the precise factors that promote TAM activation have yet to be elucidated, as each TME is characterized by unique physical and chemical conditions [13,43]. However, certain common features may be identified. For example, CSF1, IL-10 and TGF-*β* released from tumour cells and Treg cells, are powerful promoters of TAM polarization, which in turn support tumour progression by various mechanisms, like: *(i)* secretion of soluble immunosuppressive agents (IL-10, TGF-*β*, IL-1*β*), *(ii)* expression of Immune Checkpoint Inhibitors (PD-L1, B7-H4), and *(iii)* high levels of hypoxia-inducible factor 1 and 2 (HIF1, HIF2) which leads to expression of genes associated with pro-tumoural activity [12,13,43,69].

In the context of Chronic Lymphocytic Leukaemia (CLL) it has been proposed that Nurse-like cells (NLC), which are specific form of TAMs identified in this malignancy, are polarized in response to CSF-1 and HMGB1 proteins released by CLL cancer cells. In turn NLCs can stimulate and protect CLL cells by antigen presentation which stimulates BCR signaling, and also by both direct contact through membrane proteins and release of soluble factors including: [42]

- **membrane proteins:** CD2 (interacts with LFA-3 expressed on CLL cells [70]), CD31 (ligand of CD38 expressed on CLL cells), BAFF, APRIL (both BAFF and APRIL can be also released as soluble factors) [42,70]
- **soluble factors:** BDNF [71], WNT5A [72], CXCL12, CXCL13, IL6/8, IL-10 [42].

## References

1. Foey, A.D. Macrophages — Masters of Immune Activation, Suppression and Deviation. Immune Response Activation 2014. doi:10.5772/57541.

2. Martinez, F.O.; Gordon, S. The M1 and M2 paradigm of macrophage activation: Time for reassessment. F1000Prime Reports 2014, 6, 1–13.

3. Murray, P.J.; Wynn, T.A. Obstacles and opportunities for understanding macrophage polarization. J. Leukoc. Biol.2011, 89, 557–563.

4. Mantovani, A.; Sica, A.; Locati, M. Macrophage polarization comes of age. Immunity 2005, 23, 344–346. doi:10.1016/j.immuni.2005.10.001.

5. Mosser, D.M.; Edwards, J.P. Exploring the full spectrum of macrophage activation. Nature reviews immunology 2008, 8, 958–969.

6. Locati, M.; Curtale, G.; Mantovani, A. Diversity, Mechanisms, and Significance of Macrophage Plasticity. Annu. Rev. Pathol. Mech. Dis.2020, 15, 123–147. doi:10.1146/annurev-pathmechdis-012418-012718.

7. Gordon, S.; Martinez, F.O. Alternative activation of macrophages: Mechanism and functions. Immunity 2010, 32, 593–604. doi:10.1016/j.immuni.2010.05.007.

8. Ivashkiv, L.B. Epigenetic regulation of macrophage polarization and function. Trends in immunology 2013, 34, 216–223.

9. Cai, L.; Michelakos, T.; Deshpande, V.; Arora, K.S.; Yamada, T.; Ting, D.T.; Taylor, M.S.; Fernandez-del Castillo, C.; Warshaw, A.L.; Lillemoe, K.D.; others. Role of tumor-associated macrophages in the clinical course of pancreatic neuroendocrine tumors (PanNETs). Clinical Cancer Research 2019, 25, 2644–2655.

10. Grossman, J.G.; Nywening, T.M.; Belt, B.A.; Panni, R.Z.; Krasnick, B.A.; DeNardo, D.G.; Hawkins, W.G.; Goedegebuure, S.P.; Linehan, D.C.; Fields, R.C. Recruitment of CCR2+ tumor associated macrophage to sites of liver metastasis confers a poor prognosis in human colorectal cancer. Oncoimmunology 2018, 7, e1470729.

11. Wynn, T.A.; Chawla, A.; Pollard, J.W. Macrophage biology in development, homeostasis and disease. Nature 2013, 496, 445–455.

12. Bingle, L.; Brown, N.; Lewis, C.E. The role of tumour-associated macrophages in tumour progression: implications for new anticancer therapies. The Journal of Pathology: A Journal of the Pathological Society of Great Britain and Ireland 2002, 196, 254–265.

13. Mantovani, A.; Marchesi, F.; Malesci, A.; Laghi, L.; Allavena, P. Tumour-associated macrophages as treatment targets in oncology. Nature reviews Clinical oncology 2017, 14, 399.

14. DeNardo, D.G.; Brennan, D.J.; Rexhepaj, E.; Ruffell, B.; Shiao, S.L.; Madden, S.F.; Gallagher, W.M.; Wadhwani, N.; Keil, S.D.; Junaid, S.A.; others. Leukocyte complexity predicts breast cancer survival and functionally regulates response to chemotherapy. Cancer discovery 2011, 1, 54–67.

15. Doedens, A.L.; Stockmann, C.; Rubinstein, M.P.; Liao, D.; Zhang, N.; DeNardo, D.G.; Coussens, L.M.; Karin, M.; Goldrath, A.W.; Johnson, R.S. Macrophage expression of hypoxia-inducible factor-1α suppresses T-cell function and promotes tumor progression. Cancer research 2010, 70, 7465–7475.

16. Movahedi, K.; Laoui, D.; Gysemans, C.; Baeten, M.; Stangé, G.; Van den Bossche, J.; Mack, M.; Pipeleers, D.; In’t Veld, P.; De Baetselier, P.; others. Different tumor microenvironments contain functionally distinct subsets of macrophages derived from Ly6C (high) monocytes. Cancer research 2010, 70, 5728–5739.

17. Larionova, I.; Kazakova, E.; Patysheva, M.; Kzhyshkowska, J. Transcriptional, Epigenetic and Metabolic Programming of Tumor-Associated Macrophages. Cancers 2020, 12, 1411.

18. Ruffell, B.; Chang-Strachan, D.; Chan, V.; Rosenbusch, A.; Ho, C.M.; Pryer, N.; Daniel, D.; Hwang, E.S.; Rugo, H.S.; Coussens, L.M. Macrophage IL-10 blocks CD8+ T cell-dependent responses to chemotherapy by suppressing IL-12 expression in intratumoral dendritic cells. Cancer cell 2014, 26, 623–637.

19. Barabási, A.L.; Gulbahce, N.; Loscalzo, J. Network medicine: a network-based approach to human disease. Nature reviews genetics 2011, 12, 56–68.

20. Gulfidan, G.; Turanli, B.; Beklen, H.; Sinha, R.; Arga, K.Y. Pan-cancer mapping of differential protein-protein interactions. Scientific reports 2020, 10, 1–12.

21. Albert, I.; Thakar, J.; Li, S.; Zhang, R.; Albert, R. Boolean network simulations for life scientists. Source code for biology and medicine 2008, 3, 1–8.

22. Kervizic, G.; Corcos, L. Dynamical modeling of the cholesterol regulatory pathway with Boolean networks. BMC systems biology 2008, 2, 99.

23. Saadatpour, A.; Wang, R.S.; Liao, A.; Liu, X.; Loughran, T.P.; Albert, I.; Albert, R. Dynamical and structural analysis of a t cell survival network identifies novel candidate therapeutic targets for large granular lymphocyte leukemia. PLoS Computational Biology 2011, 7.

24. Naldi, A.; Carneiro, J.; Chaouiya, C.; Thieffry, D. Diversity and plasticity of Th cell types predicted from regulatory network modelling. PLoS Comput Biol 2010, 6, e1000912.

25. Alves, R.; Heiner, M.; Hiroi, N.; Chaouiya, C.; Abou-Jaoudé, W.; Traynard, P.; Monteiro, P.T.; Saez-Rodriguez, J.; Helikar, T.; Thieffry, D. Logical Modeling and Dynamical Analysis of Cellular Networks. Frontiers in Genetics www.frontiersin.org 2016, 1, 94. doi:10.3389/fgene.2016.00094.

26. Palma, A.; Jarrah, A.S.; Tieri, P.; Cesareni, G.; Castiglione, F. Gene regulatory network modeling of macrophage differentiation corroborates the continuum hypothesis of polarization states. Frontiers in physiology 2018, 9, 1659.

27. Kondratova, M.; Czerwinska, U.; Sompairac, N.; Amigorena, S.D.; Soumelis, V.; Barillot, E.; Zinovyev, A.; Kuperstein, I. A multiscale signalling network map of innate immune response in cancer reveals cell heterogeneity signatures. Nature communications 2019, 10, 1–13.

28. Razzaq, M.; Paulevé, L.; Siegel, A.; Saez-Rodriguez, J.; Bourdon, J.; Guziolowski, C. Computational discovery of dynamic cell line specific Boolean networks from multiplex time-course data. PLoS computational biology 2018, 14, e1006538.

29. Martin, S.; Zhang, Z.; Martino, A.; Faulon, J.L. Boolean dynamics of genetic regulatory networks inferred from microarray time series data. Bioinformatics 2007, 23, 866–874.

30. Maetschke, S.R.; Madhamshettiwar, P.B.; Davis, M.J.; Ragan, M.A. Supervised, semi-supervised and unsupervised inference of gene regulatory networks. Briefings in bioinformatics 2014, 15, 195–211.

31. Burger, J.A.; Tsukada, N.; Burger, M.; Zvaifler, N.J.; Dell’Aquila, M.; Kipps, T.J. Blood-derived nurse-like cells protect chronic lymphocytic leukemia B cells from spontaneous apoptosis through stromal cell–derived factor-1. Blood, The Journal of the American Society of Hematology 2000, 96, 2655–2663.

32. Tsukada, N.; Burger, J.A.; Zvaifler, N.J.; Kipps, T.J. Distinctive features of “nurselike” cells that differentiate in the context of chronic lymphocytic leukemia. Blood, The Journal of the American Society of Hematology 2002, 99, 1030–1037.

33. Boissard, F.; Fournie, J.; Quillet-Mary, A.; Ysebaert, L.; Poupot, M. Nurse-like cells mediate ibrutinib resistance in chronic lymphocytic leukemia patients. Blood cancer journal 2015, 5, e355–e355.

34. Abou-Jaoudé, W.; Traynard, P.; Monteiro, P.T.; Saez-Rodriguez, J.; Helikar, T.; Thieffry, D.; Chaouiya, C. Logical modeling and dynamical analysis of cellular networks. Frontiers in genetics 2016, 7, 94.

35. Lawrence, T.; Natoli, G. Transcriptional regulation of macrophage polarization: Enabling diversity with identity. Nature Reviews Immunology 2011, 11, 750–761. doi:10.1038/nri3088.

36. Krumsiek, J.; Marr, C.; Schroeder, T.; Theis, F.J. Hierarchical differentiation of myeloid progenitors is encoded in the transcription factor network. PLoS ONE 2011, 6.

37. G.T. Zañudo, J.; Steinway, S.N.; Albert, R. Discrete dynamic network modeling of oncogenic signaling: Mechanistic insights for personalized treatment of cancer, 2018. doi:10.1016/j.coisb.2018.02.002.

38. Emmrich, P.M.F.; Roberts, H.E.; Pancaldi, V. A Boolean gene regulatory model of heterosis and speciation. BMC evolutionary biology 2015, 15, 24.

39. Bloomingdale, P.; Niu, J.; Mager, D.E.; others. Boolean network modeling in systems pharmacology. Journal of pharmacokinetics and pharmacodynamics 2018, 45, 159–180.

40. Davidich, M.I.; Bornholdt, S. Boolean network model predicts cell cycle sequence of fission yeast. PloS one 2008, 3, e1672.

41. Orecchioni, M.; Ghosheh, Y.; Pramod, A.B.; Ley, K. Macrophage polarization: Different gene signatures in M1(Lps+) vs. Classically and M2(LPS-) vs. Alternatively activated macrophages. Frontiers in Immunology 2019, 10, 1–14.

42. en Hacken, E.; Burger, J.A. Microenvironment interactions and B-cell receptor signaling in Chronic Lymphocytic Leukemia: Implications for disease pathogenesis and treatment. Biochimica et Biophysica Acta - Molecular Cell Research 2016, 1863, 401–413. doi:10.1016/j.bbamcr.2015.07.009.

43. Guttman, O.; C. Lewis, E. M2-like macrophages and tumor-associated macrophages: overlapping and distinguishing properties en route to a safe therapeutic potential. Integrative Cancer Science and Therapeutics 2016, 3, 554–561. doi:10.15761/icst.1000204.

44. Cassetta, L.; Pollard, J.W. Targeting macrophages: therapeutic approaches in cancer. Nature Reviews Drug Discovery 2018, 17, 887–904.

45. Ogata, H.; Goto, S.; Fujibuchi, W.; Kanehisa, M. Computation with the KEGG pathway database. Biosystems 1998, 47, 119–128.

46. Zhang, B.; Srihari, S.N. Properties of Binary Vector Dissimilarity Measures. Non Journal 2000, p. 20 pp. doi:10.1117/12.473347.

47. DéBroski, R.H.; Hölscher, C.; Mohrs, M.; Arendse, B.; Schwegmann, A.; Radwanska, M.; Leeto, M.; Kirsch, R.; Hall, P.; Mossmann, H.; others. Alternative macrophage activation is essential for survival during schistosomiasis and downmodulates T helper 1 responses and immunopathology. Immunity 2004, 20, 623–635.

48. Müssel, C.; Hopfensitz, M.; Kestler, H.A. BoolNet—an R package for generation, reconstruction and analysis of Boolean networks. Bioinformatics 2010, 26, 1378–1380.

49. Chawla, A. Control of macrophage activation and function by PPARs. Circulation research 2010, 106, 1559–1569.

50. Murray, P.J. Macrophage polarization. Annual review of physiology 2017, 79, 541–566.

51. Rutschman, R.; Lang, R.; Hesse, M.; Ihle, J.N.; Wynn, T.A.; Murray, P.J. Cutting edge: Stat6-dependent substrate depletion regulates nitric oxide production. The Journal of Immunology 2001, 166, 2173–2177.

52. Vannella, K.M.; Barron, L.; Borthwick, L.A.; Kindrachuk, K.N.; Narasimhan, P.B.; Hart, K.M.; Thompson, R.W.; White, S.; Cheever, A.W.; Ramalingam, T.R.; others. Incomplete deletion of IL-4Rα by LysM Cre reveals distinct subsets of M2 macrophages controlling inflammation and fibrosis in chronic schistosomiasis. PLoS Pathog 2014, 10, e1004372.

53. Dalmas, E.; Toubal, A.; Alzaid, F.; Blazek, K.; Eames, H.L.; Lebozec, K.; Pini, M.; Hainault, I.; Montastier, E.; Denis, R.G.; others. Irf5 deficiency in macrophages promotes beneficial adipose tissue expansion and insulin sensitivity during obesity. Nature medicine 2015, 21, 610–618.

54. Friedrich, J.; Heim, L.; Trufa, D.I.; Sirbu, H.; Rieker, R.J.; Chiriac, M.T.; Finotto, S. STAT1 deficiency supports PD-1/PD-L1 signaling resulting in dysfunctional TNFα mediated immune responses in a model of NSCLC. Oncotarget 2018, 9, 37157.

55. Satoh, T.; Takeuchi, O.; Vandenbon, A.; Yasuda, K.; Tanaka, Y.; Kumagai, Y.; Miyake, T.; Matsushita, K.; Okazaki, T.; Saitoh, T.; others. The Jmjd3-Irf4 axis regulates M2 macrophage polarization and host responses against helminth infection. Nature immunology 2010, 11, 936–944.

56. Murtagh, F.; Contreras, P. Algorithms for hierarchical clustering: an overview. Wiley Interdisciplinary Reviews: Data Mining and Knowledge Discovery 2012, 2, 86–97.

57. McInnes, L.; Healy, J.; Astels, S. hdbscan: Hierarchical density based clustering. Journal of Open Source Software 2017, 2, 205.

58. Eduati, F.; De Las Rivas, J.; Di Camillo, B.; Toffolo, G.; Saez-Rodriguez, J. Integrating literature-constrained and data-driven inference of signalling networks. Bioinformatics 2012, 28, 2311–2317.

59. Kulkarni, S.R.; Vandepoele, K. Inference of plant gene regulatory networks using data-driven methods: A practical overview. Biochimica et Biophysica Acta (BBA)-Gene Regulatory Mechanisms 2020, 1863, 194447.

60. Macklin, P.; Frieboes, H.B.; Sparks, J.L.; Ghaffarizadeh, A.; Friedman, S.H.; Juarez, E.F.; Jonckheere, E.; Mumenthaler, S.M. Progress towards computational 3-d multicellular systems biology. In Systems Biology of Tumor Microenvironment; Springer, 2016; pp. 225–246.

61. Margaris, K.; Black, R.A. Modelling the lymphatic system: challenges and opportunities. Journal of the Royal Society Interface 2012, 9, 601–612.

62. Galle, J.; Aust, G.; Schaller, G.; Beyer, T.; Drasdo, D. Individual cell-based models of the spatial-temporal organization of multicellular systems—Achievements and limitations. Cytometry Part A: The Journal of the International Society for Analytical Cytology 2006, 69, 704–710.

63. Albert, R.; Robeva, R. Signaling networks: Asynchronous boolean models. In Algebraic and discrete mathematical methods for modern biology; Elsevier, 2015; pp. 65–91.

64. Ihaka, R.; Gentleman, R. R: a language for data analysis and graphics. Journal of computational and graphical statistics 1996, 5, 299–314.

65. Holland, C.H.; Valdeolivas, A.; Saez-Rodriguez, J. TF activity inference from bulk transcriptomic data with DoRothEA as regulon resource.

66. Garcia-Alonso, L.; Holland, C.H.; Ibrahim, M.M.; Turei, D.; Saez-Rodriguez, J. Benchmark and integration of resources for the estimation of human transcription factor activities. Genome research 2019, 29, 1363–1375.

67. Geeraerts, X.; Bolli, E.; Fendt, S.M.; Van Ginderachter, J.A. Macrophage metabolism as therapeutic target for cancer, atherosclerosis, and obesity. Frontiers in Immunology 2017, 8, 289.

68. Wang, Z.; Brandt, S.; Medeiros, A.; Wang, S.; Wu, H.; Dent, A.; Serezani, C.H. MicroRNA 21 is a homeostatic regulator of macrophage polarization and prevents prostaglandin E 2-mediated M2 generation. PloS one 2015, 10, e0115855.

69. Heusinkveld, M.; van der Burg, S.H. Identification and manipulation of tumor associated macrophages in human cancers. Journal of translational medicine 2011, 9, 216.

70. Boissard, F.; Tosolini, M.; Ligat, L.; Quillet-Mary, A.; Lopez, F.; Fournié, J.J.; Ysebaert, L.; Poupot, M. Nurse-like cells promote CLL survival through LFA-3/CD2 interactions. Oncotarget 2017, 8, 52225.

71. Talbot, H.; Saada, S.; Barthout, E.; Gallet, P.F.; Gachard, N.; Abraham, J.; Jaccard, A.; Troutaud, D.; Lalloué, F.; Naves, T.; others. BDNF belongs to the nurse-like cell secretome and supports survival of B chronic lymphocytic leukemia cells. Scientific reports 2020, 10, 1–9.

72. Kipps, T.J.; Stevenson, F.K.; Wu, C.J.; Croce, C.M.; Packham, G.; Wierda, W.G.; O’brien, S.; Gribben, J.; Rai, K. Chronic lymphocytic leukaemia. Nature reviews Disease primers 2017, 3, 1–22.

